## Supplementary file for "Neutralizing gut-derived lipopolysaccharide as a novel therapeutic strategy for severe leptospirosis"

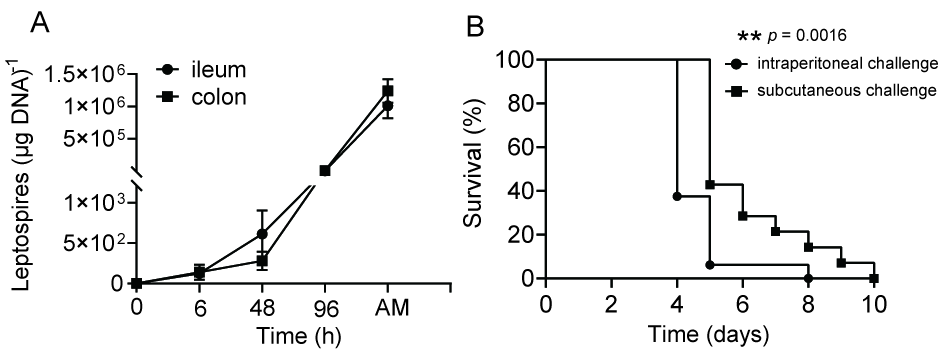

**Supplementary Fig. S1**. **The effect of challenge routes on leptospirosis**. Six-week-old female hamsters were injected intraperitoneally or subcutaneously with 10^7^ leptospires. Then, hamsters were treated as described in the Methods. (A) Leptospiral load in the ileums and colons of hamsters (n = 4) infected subcutaneously were determined by qPCR. (B) Survival curves of hamsters infected intraperitoneally (n = 16) or subcutaneously (n = 14) with 10^7^ leptospires. Each experiment was repeated three times. Data are shown as the mean ± SEM and analyzed by the Wilcoxon rank-sum test. Survival differences was analyzed by the Kaplan-Meier log rank test. ***p* < 0.01.

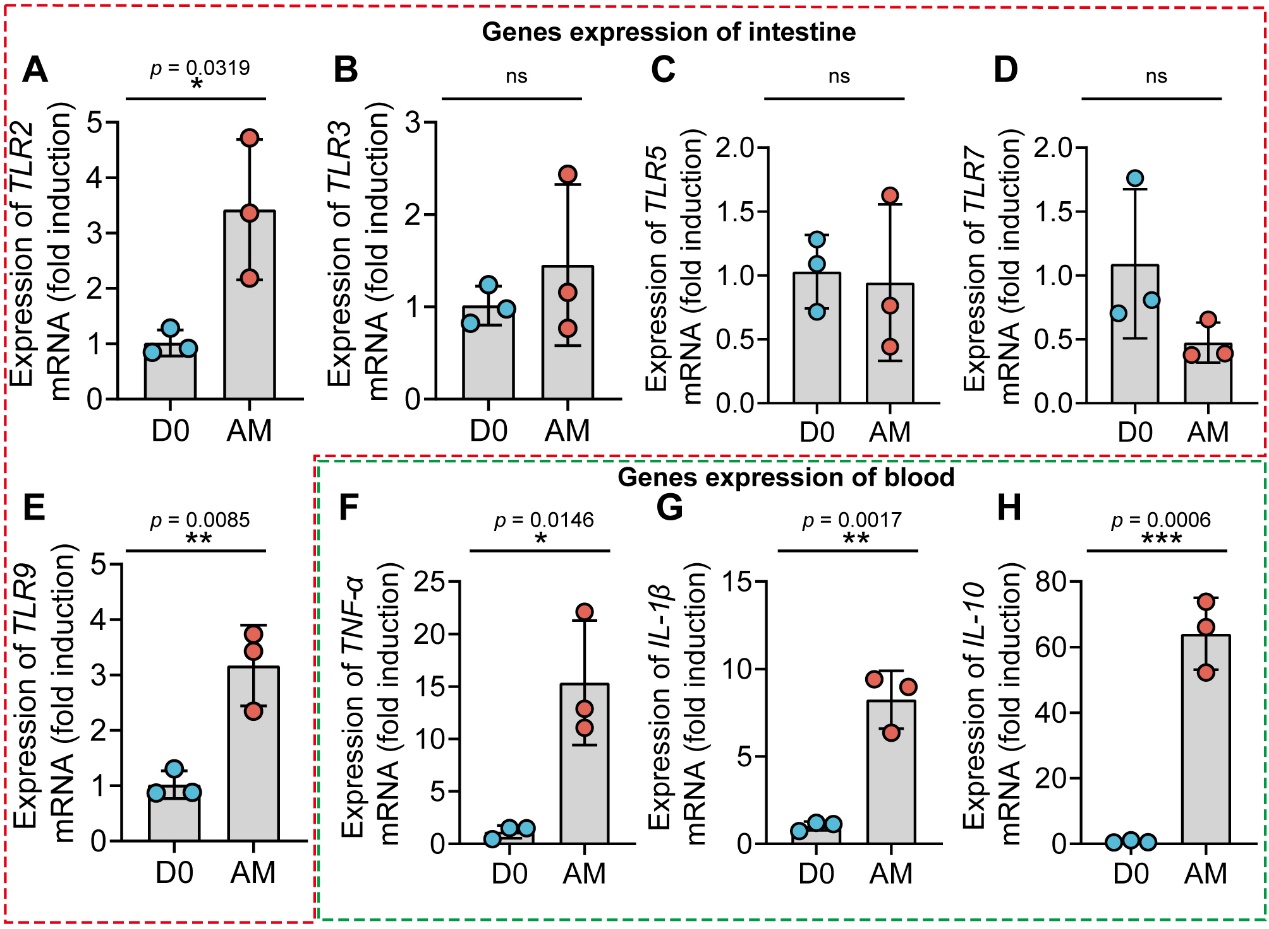

**Supplementary Fig. S2. The gene expression of TLRs in the intestine and inflammatory cytokines in the blood.** Six-week-old female hamsters were injected intraperitoneally with 10^7^ leptospires. Hamsters were euthanatized at 0 h and AM p.i.. Colons and blood were collected aseptically for analysis of gene expression. D0, uninfected hamster. AM, articulo mortis. (A-E) The gene expression of *TLR2* (A), *TLR3* (B), *TLR5* (C), *TLR7* (D), and *TLR9* (E) in the intestine was analyzed by RT-qPCR. (F-H) The gene expression of *TNF-α* (F), *IL-1β* (G), and *IL-10* (H) in the blood was analyzed by RT-qPCR. The mRNA levels of genes of D0 group (n = 3) and AM group (n = 3) were normalized to the expression of the housekeeping gene *GAPDH*. The mRNA levels of cytokines in uninfected controls were set as 1.0. Each infection experiment was repeated three times. Data are shown as the mean ± SEM and analyzed by the Wilcoxon rank-sum test. **p* < 0.05, ***p* < 0.01, ****p* < 0.001. ns, not significant.

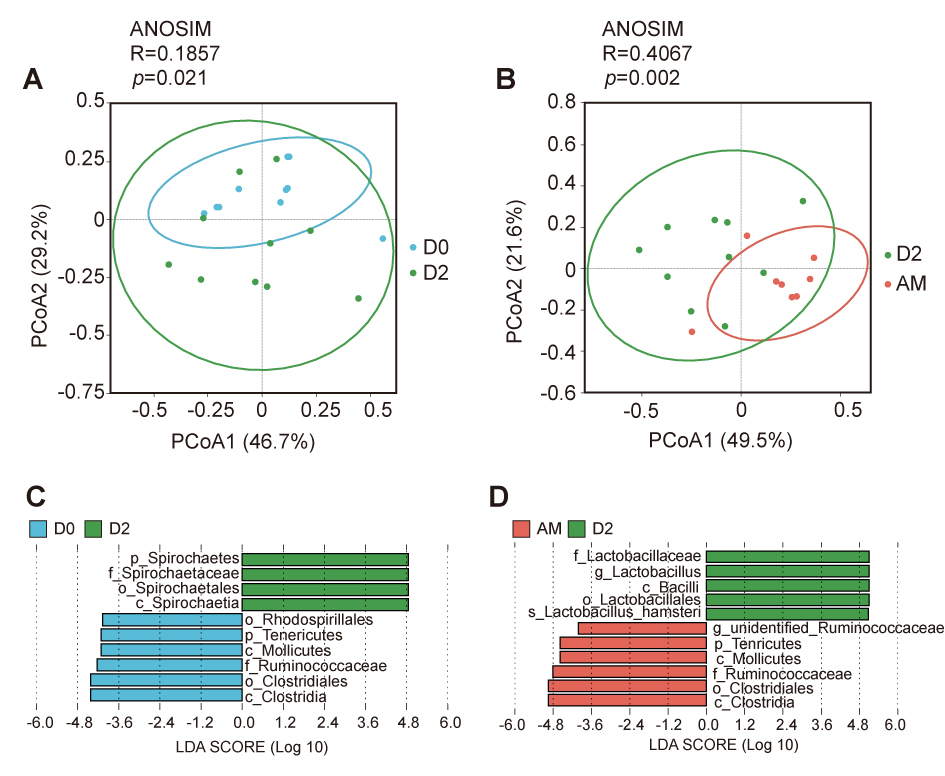

**Supplementary Fig. S3**. **The effect of *Leptospira* infection on the composition of the gut microbiota.** (A) PCoA of fecal samples based on 16S rRNA gene sequencing using weighted UniFrac (D0, n = 10; D2, n = 10). (B) PCoA of fecal samples based on 16S rRNA gene sequencing using weighted UniFrac (D2, n = 10; AM, n = 8). (C) LEfSe between D0 and D2 hamsters (LDA score > 4). Cambridge blue bars indicate taxa enrichment in D0 hamsters, and green bars indicate taxa enrichment in D2 hamsters. (D) LEfSe between D2 and AM hamsters (LDA score > 4). Green bars indicate taxa enrichment in D2 hamsters, and pink bars indicate taxa enrichment in AM hamsters.

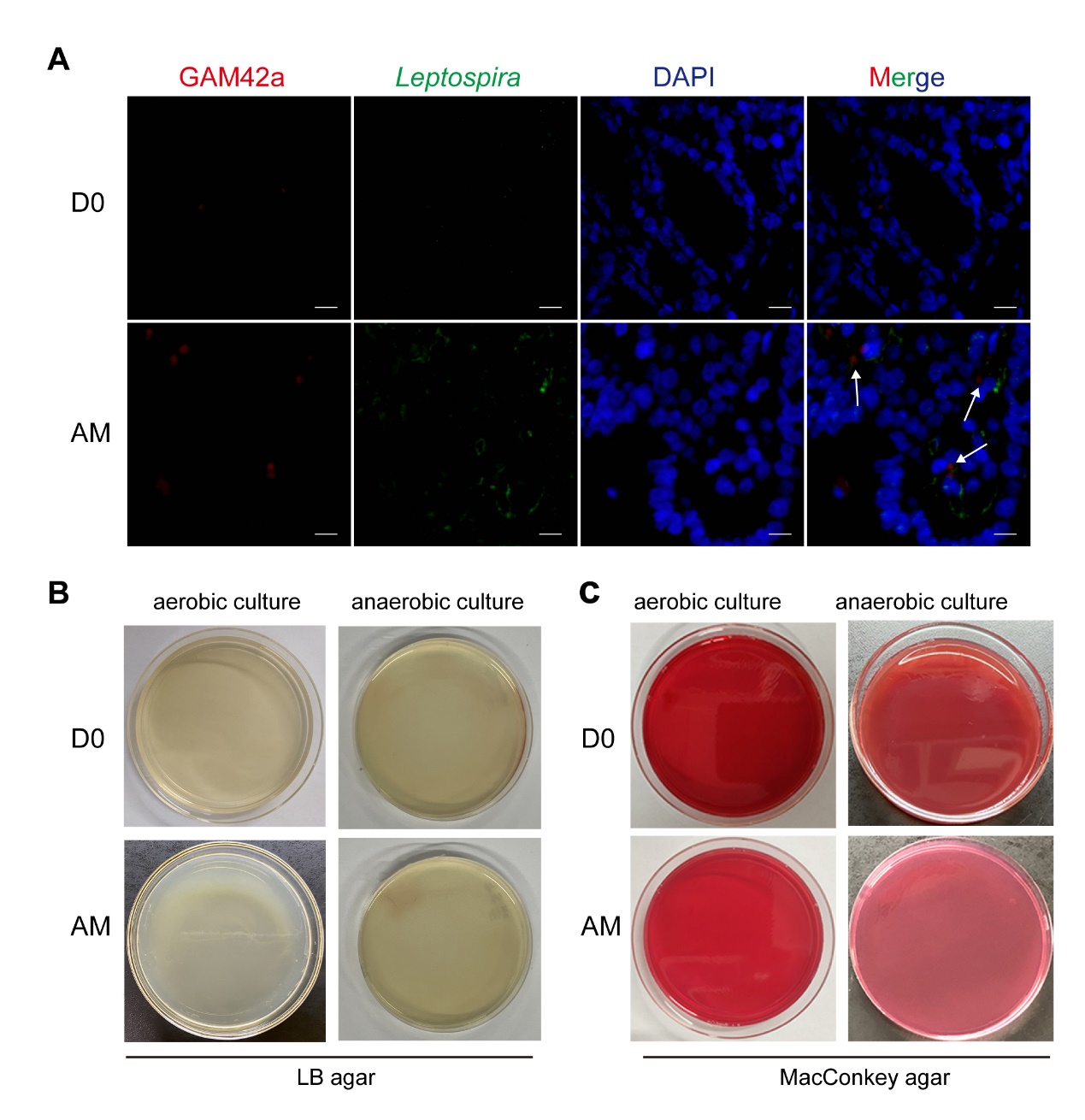

**Supplementary Fig. S4**. **Translocation of the Proteobacteria during *L. interrogans* infection.** Six-week-old female hamsters were intraperitoneally infected with 10^7^ leptospires. Colons of the D0 group and the AM group were collected and detected by FISH and IF. Blood from the D0 group and the AM group were plated on LB agar or MacConkey agar plates aerobically or anaerobically at 37°C for 36 h. D0, uninfected hamster. AM, articulo mortis. (A) Colons of D0 (n = 4) and AM (n = 4) were examined by FISH with GAM42a probe (Red) and by IF with anti-*Leptospira* serum (green). Scale bar, 20 μm. (B-C) The blood of the D0 (n = 4) hamsters and AM hamsters (n = 4) were aerobically or anaerobically cultured on LB plates (B) or MacConkey plates (C). Each experiment was repeated three times.

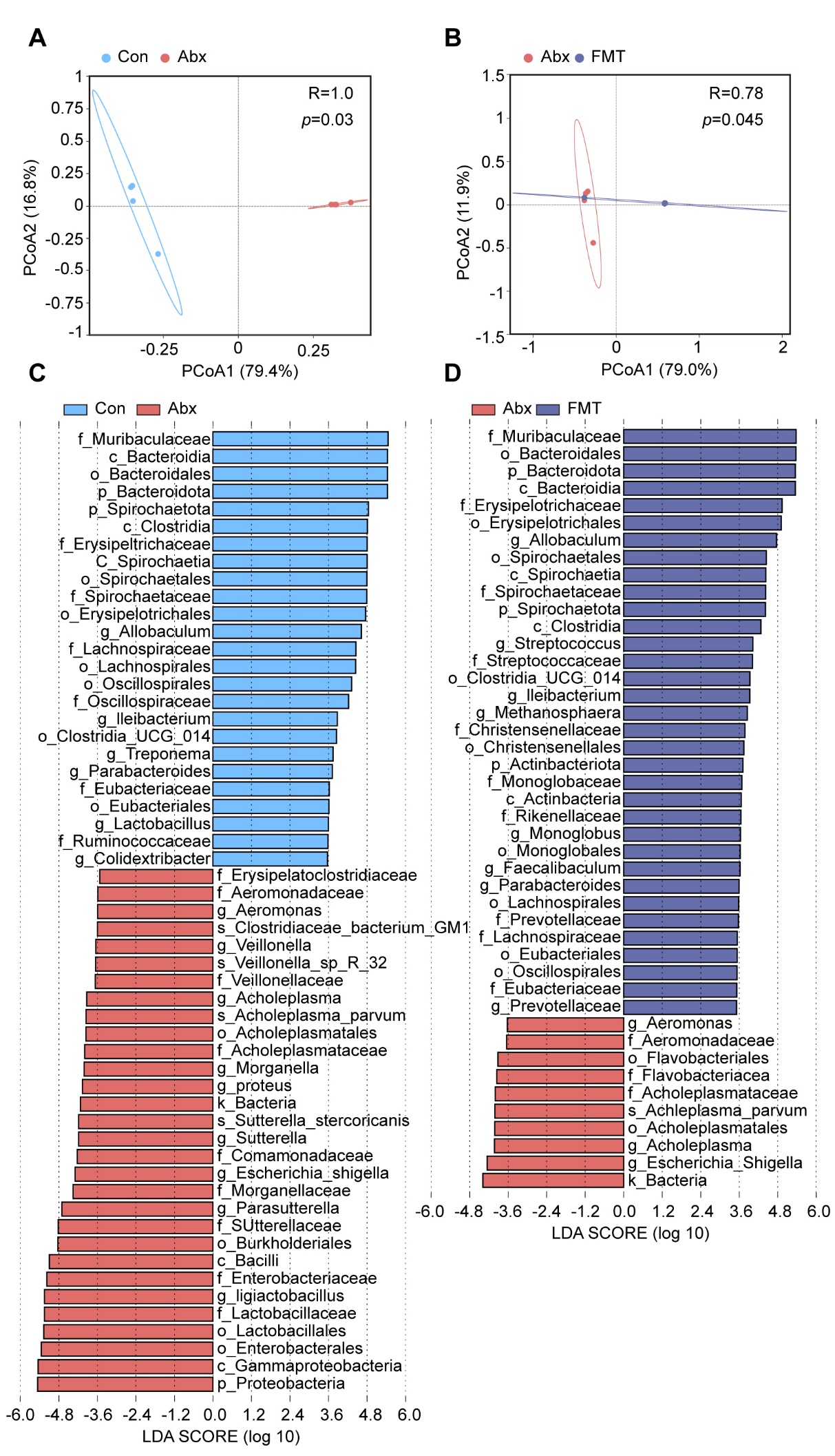

**Supplementary Fig. S5. The effect of Abx treatment on the composition of the gut microbiota.** (A) PCoA of fecal samples based on 16S rRNA gene sequencing using weighted UniFrac (Con, n = 4; Abx, n = 4). (B) PCoA of fecal samples based on 16S rRNA gene sequencing using weighted UniFrac (Abx, n = 4; FMT, n = 4). (C) LEfSe between Con and Abx hamsters (LDA score > 4). Cambridge blue bars indicate taxa enrichment in Con hamsters, and pink bars indicate taxa enrichment in Abx hamsters. (D) LEfSe between FMT and Abx hamsters (LDA score > 4). Blue bars indicate taxa enrichment in FMT hamsters, and pink bars indicate taxa enrichment in Abx hamsters.

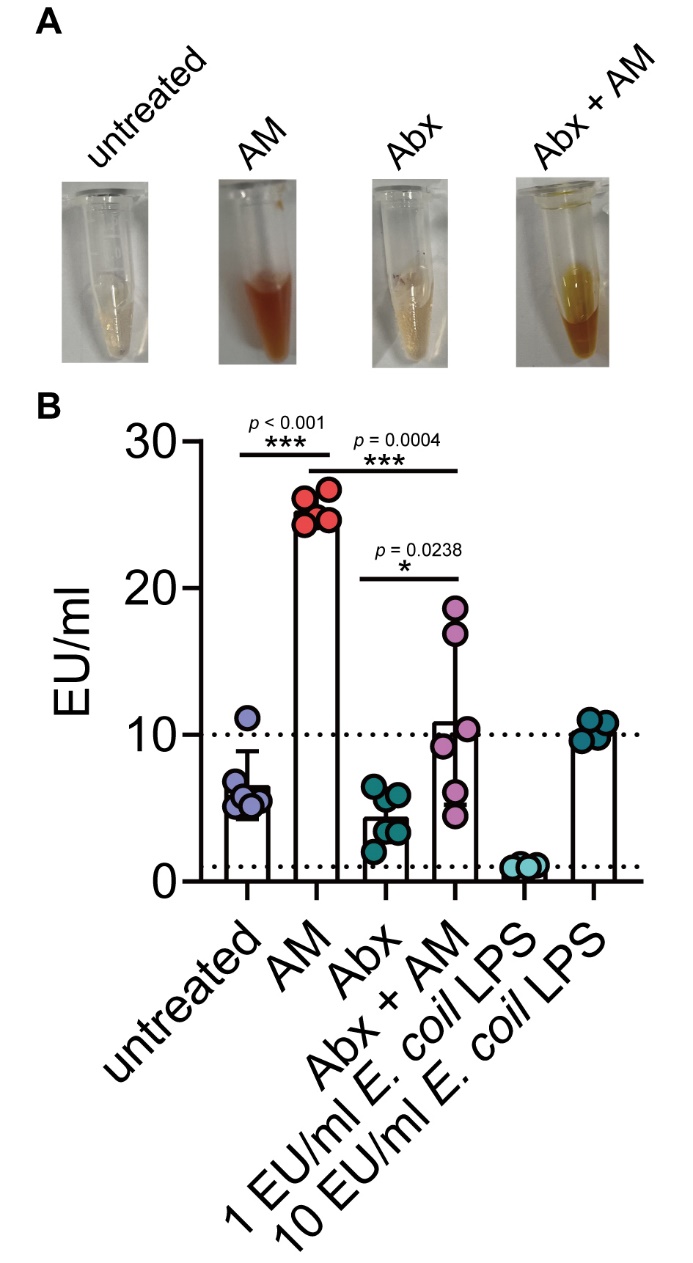

**Supplementary Fig. S6**. ***Leptospira* infection increased LPS level in the blood.** Six-week-old female hamsters were intraperitoneally infected with 10^7^ leptospires. The serum of untreated hamsters (n = 6), Abx-treated hamsters (n = 6), AM hamsters (n = 5), and Abx-treated AM hamsters (n = 6) were collected for LPS detection. *E. coil* LPS were used as a positive control to examine the quality of the kits. (A) Representative pictures of collected serum from untreated group, Abx-treated group, AM group, and Abx-treated AM group. (B) LPS level in the untreated group, Abx-treated group, AM group, Abx-treated AM group, 1 EU/ml *E. coil* LPS group, and 10 EU/ml *E. coil* LPS group. Each experiment was repeated three times. Data are shown as the mean ± SEM and analyzed by the Wilcoxon rank-sum test. **p* < 0.05, ****p* < 0.001.

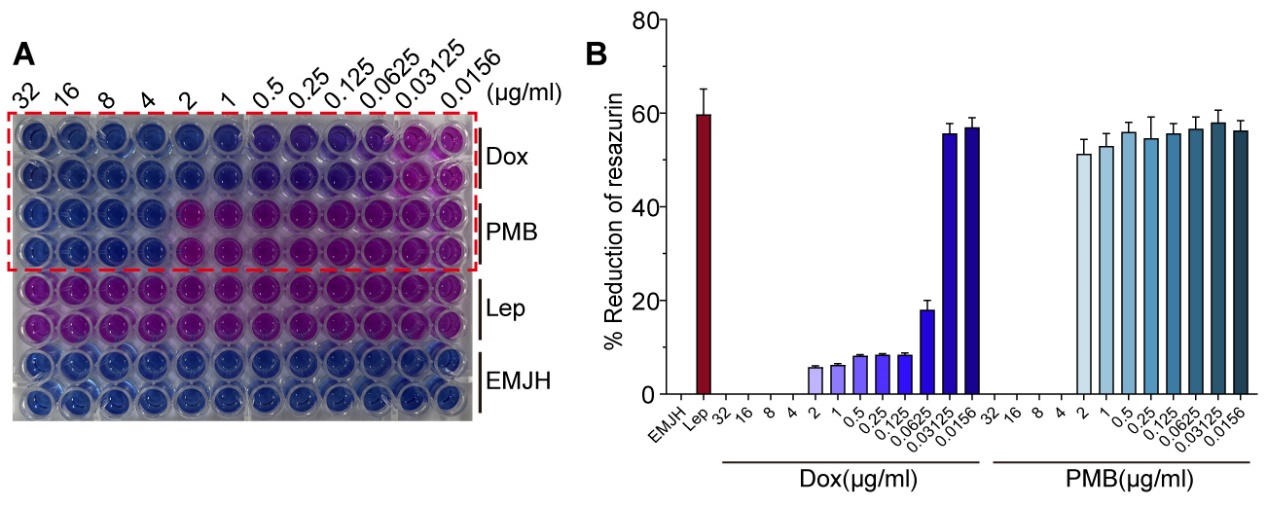

**Supplementary Fig. S7**. **Determination minimum inhibitory concentrations of PMB and Dox on *Leptospira* 56601**. Leptospires at exponential-phase were deposited at a final concentration of 2×10^6^ leptospires/mL in each well of 96 well plates with serial two-fold dilutions of PMB and Dox ranging from 32 to 0.0156 μg/L in EMJH. The final volume in each well was 200 μL. The plates were incubated for 3 days at 30 °C, then 20 μL of Alamar Blue was added to each well and the samples were incubated at 30 °C for 2 days. Each strain–drug combination was tested in duplicate, and positive (bacteria and no antibiotic added) and negative (no bacteria added) controls were included in each plate. (A) Representative pictures of Alamar Blue test. (B) The percentage of resazurin reduction at different dosage of PMB and Dox incubated with leptospires. Each experiment was repeated three times. Data are shown as the mean ± SEM.

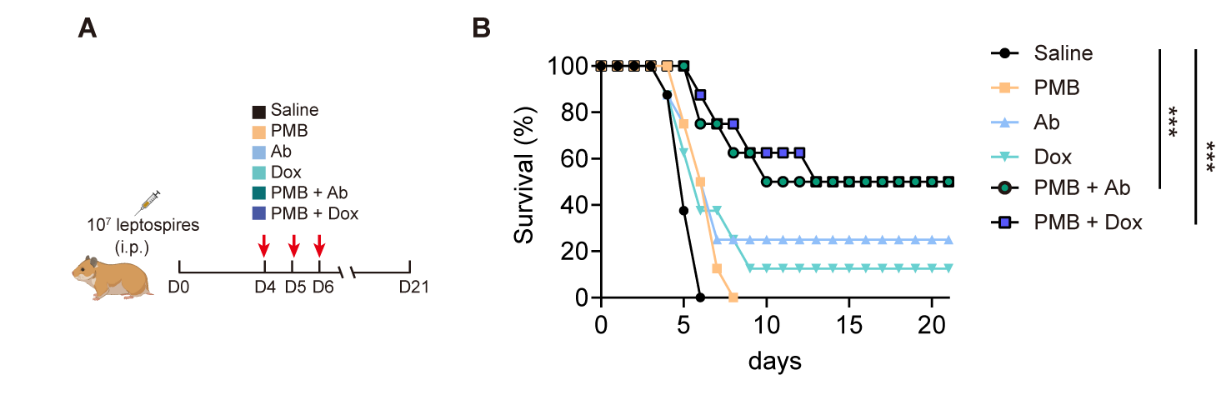

**Supplementary Fig. S8. LPS neutralization combined with antibody therapy or antibiotic therapy improved the survival rate in male hamsters of severe leptospirosis.** (A) Flow diagram of the experiment. Six-week-old male hamsters were injected intraperitoneally with 10^7^ leptospires. Group 1: saline control; Group 2: Polymyxin B (PMB) (1 mg/kg, i.p.); Group 3: Antibody against *Leptospira* (Ab) (16 mg/kg, subcutaneous injection); Group 4: Doxycycline (Dox) (5 mg/kg, i.p.); Group 5: PMB (1 mg/kg, i.p.) and Ab (16 mg/kg, subcutaneous injection); Group 6: PMB (1 mg/kg, i.p.) and Dox (5 mg/kg, i.p.). Hamsters were treated on 3 consecutive days, twice a day. (B) Survival rate of male hamsters of severe leptospirosis after different treatments Hamsters were monitored daily for 21 days. n = 8 per group. Each infection experiment was repeated three times. Survival differences between the study groups were compared by using the Kaplan-Meier log-rank test. **p* < 0.05, ****p* < 0.001.

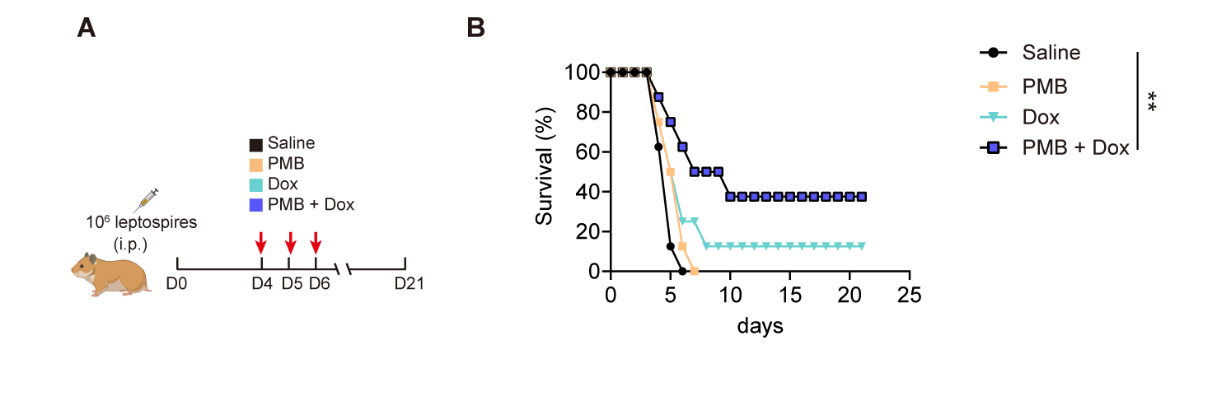

**Supplementary Fig. S9. LPS neutralization combined with antibiotic therapy improved the survival rate in female hamsters infected by 56606.** (A) Flow diagram of the experiment. Six-week-old female hamsters were injected intraperitoneally with 10^6^ leptospires (56606). Group 1: saline control; Group 2: Polymyxin B (PMB) (1 mg/kg, i.p.); Group 3: Doxycycline (Dox) (5 mg/kg, i.p.); Group 4: PMB (1 mg/kg, i.p.) and Ab (16 mg/kg, subcutaneous injection); Hamsters were treated on 3 consecutive days, twice a day. Hamsters were monitored daily for 21 days. (B) Survival rate of female hamsters of severe leptospirosis after different treatments. n = 8 per group. Each infection experiment was repeated three times. Survival differences between the study groups were compared by using the Kaplan-Meier log-rank test. ***p* < 0.01.
