## Supplementary material for "Neutralizing gut-derived lipopolysaccharide as a novel therapeutic strategy for severe leptospirosis": Table S1

| Gene | Primer type | Sequence (5’-3’) | Identifier |
| --- | --- | --- | --- |
| Hamster GAPDH | Sense | GATGCTGGTGCCGAGTATGT | [1] |
|  | Antisense | GCAGAAGGTGCGGAGATGA |  |
| Hamster Claudin-2 | Sense | ACTGTCCACGGCTACCAACTACT | This study |
|  | Antisense | TCTGGCAGGGAATGTTGGAG |  |
| Hamster Claudin-3 | Sense | GCTCACCTTAGTACCCGTGTCCT | This study |
|  | Antisense | GGCAGGAGCAGCAGAGCAA |  |
| Hamster Claudin-4 | Sense | GGGGATGCTTCTCTCAGTGGT | This study |
|  | Antisense | GACACAGGCACCATGGCC |  |
| Hamster Zo-1 | Sense | GGAGAGGTGTTCCGTGTTGTG | This study |
|  | Antisense | ACTGCTCAGCTCTGTTCTTATTGG |  |
| Hamster JAMA | Sense | TGACCTGCTCCGAACAAGATG | This study |
|  | Antisense | ACTTTGGATCAACCGTGTATGAA |  |
| Hamster Mucin-2 | Sense | ATCACTTTCCAGGCCATTGAAG | This study |
|  | Antisense | CTACAGGTTCAGTTGGTCCCAGT |  |
| Hamster TNF-α | Sense | GGTGATACCAGCAGACGG | [1] |
|  | Antisense | CTTGATGGCGGACAGGA |  |
| Hamster IL-1β | Sense | TTCTGTGACTCCTGGGATGGT | [2] |
|  | Antisense | GTTGGTTTATGTTCTGTCCGTTG |  |
| Hamster IL-10 | Sense | AAGGGTTACTTGGGTTGCC | [1] |
|  | Antisense | AATGCTCCTTGATTTCTGGC |  |
| Hamster TLR9 | Sense | AGGTAAAGTGTGGCAGTCCCG | This study |
|  | Antisense | CCAAAACAATCCCAGGAAAGG |  |
| Hamster TLR7 | Sense | TTCTCCCCAACCTTGTCCAGT | This study |
|  | Antisense | ATGAAGTTAGTGCCAAGGTCAAGAA |  |
| Hamster TLR5 | Sense | GGGTCCCTGTCCCAGTATCAA | This study |
|  | Antisense | TGCCCGGAGAGTTTATTGAGAA |  |
| Hamster TLR3 | Sense | AAGCATTGCCTGGTTCGTTAG | This study |
|  | Antisense | GTATCAAACAGCATCACTGGGAA |  |
| Hamster TLR2 | Sense | TGTTTCCCGTGTTACTGGTCAT | [1] |
|  | Antisense | CACCTGCTTCCAGACTCACC |  |
| Hamster TLR4 | Sense | ACGACGAGGACTGGGTGAGA | [1] |
|  | Antisense | GCCTTCCTGGATGATGTTGG |  |
| GAM42a (Gammaproteobacteria) | probe | GCCTTCCCACATCGTTT | [3] |
| EUB338 (universal) | probe | GCTGCCTCCCGTAGGAGT | [4] |
