## Supplementary material for "Neutralizing gut-derived lipopolysaccharide as a novel therapeutic strategy for severe leptospirosis": Table S2

| Table 2 Histological colitis scoring system | | |
| --- | --- | --- |
| Feature score | Score | Description |
| Inflammation severity | 0 | None |
|  | 1 | Minimal in mucosa |
|  | 2 | Mild affecting mucosa and sub-mucosa |
|  | 3 | Moderate affecting mucosa, sub-mucosa and sometimes transmural |
|  | 4 | Severe: often transmural |
| Erosion and ulceration | 0 | None |
|  | 1 | Rare erosions |
|  | 2 | Some erosions |
|  | 3 | Multiple erosions and ulcerations with cryptitis and crypt abscesses |
|  | 4 | Ulcers associated with necrosis and fibrosis |
| Epithelial hyperplasia | 0 | None |
|  | 1 | Mild |
|  | 2 | Mild with minimum goblet cell loss |
|  | 3 | Moderate with mild goblet cell loss |
|  | 4 | Marked with moderate to marked goblet cell loss |
| Per cent involvement | 0 | less 10% |
|  | 1 | 1-25% |
|  | 2 | 26-50% |
|  | 3 | 51-75% |
|  | 4 | 76-100% |
